# Calcium Sulfate Microparticle Size Modification for Improved Alginate Hydrogel Fabrication and Its Application in 3D Cell Culture

**DOI:** 10.1101/2023.10.05.560969

**Authors:** Joo Ho Kim, Siddharth Iyer, Christian Tessman, Shashank Vummidi Lakshman, Heemin Kang, Luo Gu

**Author notes:** These authors contributed equally to this work (Joo Ho Kim, Siddharth Iyer).

## Abstract

Calcium ion-crosslinked alginate hydrogels are widely used as a materials system for investigating cell behavior in 3D environments *in vitro*. Suspensions of calcium sulfate particles are often used as the source of Ca^2+^ to control the rate of gelation. However, the instability of calcium sulfate suspensions can increase chances of reduced homogeneity of the resulting gel and requires researcher’s proficiency. Here, we show that ball-milled calcium sulfate microparticles with smaller sizes can create more stable crosslinker suspensions than unprocessed or simply autoclaved calcium sulfate particles. In particular, 15 µm ball-milled calcium sulfate microparticles result in gels that are more homogeneous with a balanced gelation rate, which facilitates fabrication of gels with consistent mechanical properties and reliable performance for 3D cell culture. Overall, these microparticles represent an improved method for alginate hydrogel fabrication that can increase experimental reliability and quality for 3D cell culture.

## 1. Introduction

The importance of substrate properties on cell phenotype has been widely studied over the last few decades. Researchers have discovered that cells can sense and respond to the stiffness cues from their microenvironment [1–7]. In addition to stiffness, matrix viscoelasticity (e.g. stress-relaxation or creep) is identified as an important property that regulates cellular behaviors [8–13]. In turn, this knowledge has enabled us to fine tune *in vitro* 3D cell culture for the purpose of tissue engineering, so that one can mimic different mechanical factors of extracellular matrix (ECM) to control cell spreading, proliferation, and differentiation, etc. Specifically, alginate hydrogels with tunable viscoelasticity have recently been developed as a powerful tool for mimicking physiologically relevant viscoelastic microenvironments and controlling cell phenotype [8, 9, 14–16].

Alginate is a polysaccharide derived from seaweed, and it has a copolymer structure composed of repeating units of guluronic acid and mannuronic acid. Alginate can form hydrogels by crosslinking blocks of repeating guluronic acid residues via addition of divalent ions such as Ca^2+^. Ionically crosslinked alginate hydrogels are desirable as substrates for 3D cell culture due to their biocompatibility, mechanical tunability, and mild, cell-friendly gelation process [17]. We have previously shown that low molecular weight (MW) alginate crosslinked with Ca^2+^ results in gels that exhibited faster stress relaxation due to higher chain mobility; Conversely, high molecular weight alginate results in gels with slower stress relaxation [9]. This property allows for the independent variation of gel stress relaxation and stiffness and the creation of controlled viscoelastic microenvironments that have provided key insights into stem cell transcriptional regulation [18].

Viscoelastic alginate hydrogels for 3D cell culture commonly use suspensions of autoclave sterilized calcium sulfate dihydrate (CaSO_4_ꞏ2H_2_O) particles for ionic crosslinking [9, 18]. Compared to calcium chloride or similar calcium salts, calcium sulfate dihydrate (calcium sulfate from here on) has low solubility in water and forms a particulate suspension. These suspended particles delay the gelation process (delayed release of Ca^2+^) where the cells and alginate polymer solution can be well mixed with the calcium sulfate suspension and then molded into a desired shape before the gel fully solidifies (Fig. 1a). However, low suspension stability of large calcium sulfate particles can increase the variability and local heterogeneity of the hydrogels.

**Figure 1.**
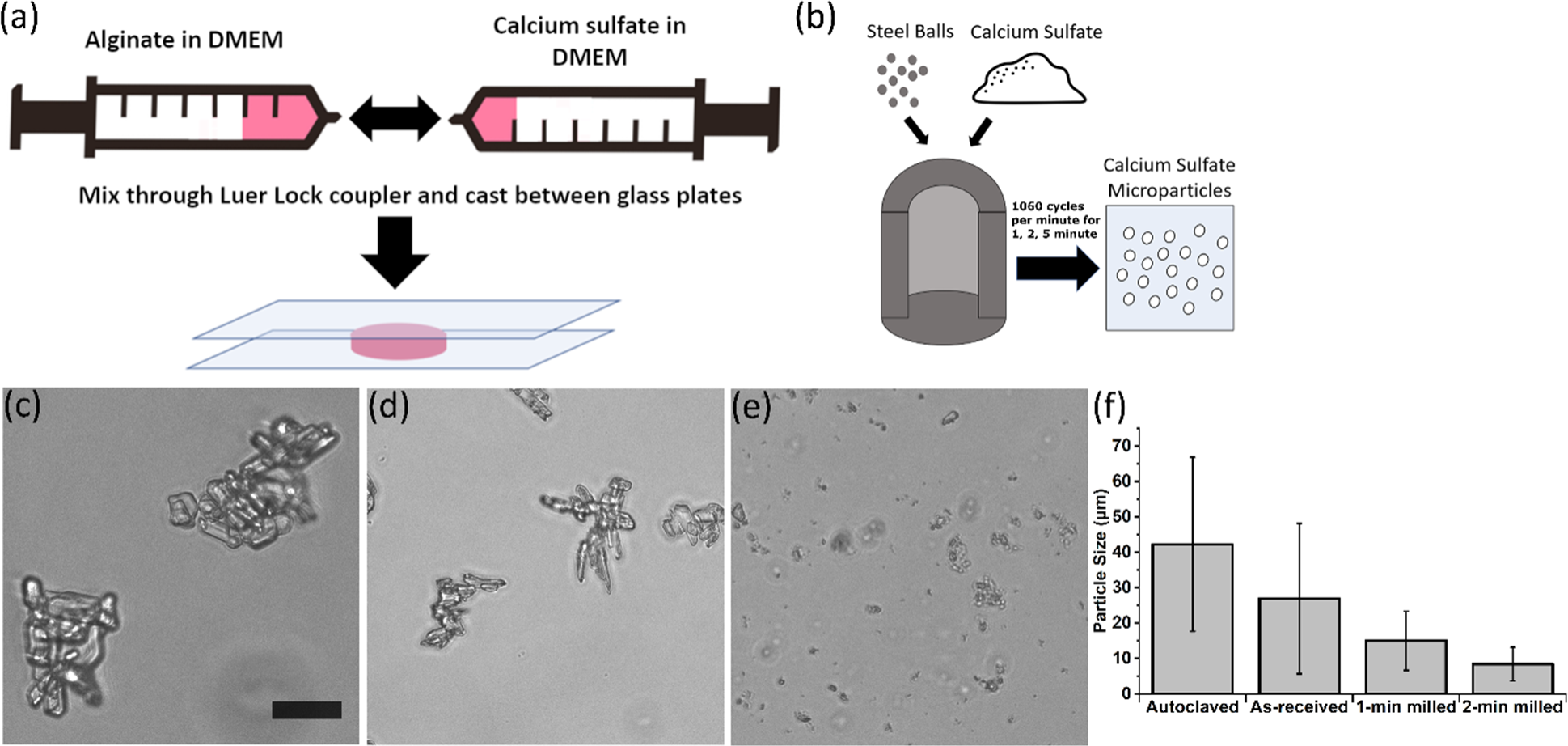
The effect of ball milling treatment on particle size and morphology. (a) General depiction of alginate hydrogel fabrication method. Two different syringes containing 1) alginate (with or without cells) and 2) calcium sulfate suspension are connected with Luerlock coupler and mixed. Mixed solution is then casted between glass plates with spacers to mold into plate-like shape. Cylindrical disks are then punched out using biopsy punches for mechanical testing and 3D cell culture experiments. (b) Depiction of ball-milling method. (c) Optical microscopy image of autoclaved particles. (d) Optical microscopy image of as-received particles. (e) Optical microscopy image of 1-minute milled particles. Optical image scale bar represents 50 μm. (f) Average particle size results from optical microscopy (n=100 per condition; error bars represent standard deviation).

Here, we report a mechanical ball milling method to modify calcium sulfate microparticles, which generated microparticles with different sizes. By examining the effect of particle size on suspension stability and hydrogel fabrication, we found calcium sulfate particles with an average size ∼15 µm form more stable suspensions while enabling reliable fabrication for homogeneous viscoelastic hydrogels with both high and low molecular weight alginates.

## 2. Methods

### 2.1 Calcium Sulfate Milling and Powder Preparation

For autoclaved particles, 8.4 g of calcium sulfate dihydrate (Acros Organics; Cat# 22527500) was suspended into 40 mL of deionized (DI) water and then autoclave sterilized. To make a working solution for gel fabrication, this autoclaved stock solution was diluted at a 4:6 ratio into Dulbecco’s Modified Eagle Medium (DMEM) to obtain 0.084g/mL working solution.

For milled particles, calcium sulfate was milled using a shaker mill (SPEX 8000D Shaker Mill) with 5 mm diameter stainless steel balls at 1060 cycles per minute for 1, 2, or 5 minutes with ball-to-powder mass ratio of 5 and total mass of 9.6 g. After washing twice with 50 mL of 70% ethanol (EtOH), the milled particles were sterilized in 70% EtOH for 48 hours. Then, the sterilized particles were centrifuged at 300 x g for 5 minutes to remove EtOH and replace it with sterile DI water. After vortexing, the particles were snap-frozen using liquid nitrogen and lyophilized for 96 hours. 10 mL of DMEM was added to 0.84 g of particles (milled or as-received after ethanol sterilization) to create a working solution for gel fabrication. For the milled particles, ultra-fine particles were removed from the working solution by vortexing for 1 minute and removing 5 mL of DMEM supernatant once supernatant settled without distinct visible particles. The concentration was then gravimetrically measured and adjusted back to 0.084 g/mL.

### 2.2 Imaging

Inverted optical microscope (EVOS M5000) with 20x objective was used to obtain images of particles. Calcium sulfate particles were suspended in water with 0.05% Triton X-100 as a surfactant to disaggregate particle clumps and measured using ImageJ. The length of the longest axis of a single particle was measured as the representative particle size. For each condition, 100 particles were measured to obtain average size of calcium sulfate particles.

### 2.3 Suspension Stability

To examine the suspension stability, working solution was prepared for each calcium sulfate condition according to the method described above. For each condition, the working solution was mixed continuously in a 50 mL conical tube. While mixing, 2 mL of the solution was pipetted from ∼1/5 height of the conical tube and pipetted into a glass cuvette. After 10 sec, 30 sec, 60 sec, or 120 sec of settling had occurred, 200 μL of working solution was removed from the top of the solution in the cuvette. The 200 μL of suspension was pipetted into a microtube for further concentration measurement. For each measurement, new suspension was prepared and used only once.

The suspension containing microtubes were then frozen and lyophilized for 48 hours, leaving dry calcium sulfate particles. This was repeated for each time period of settling. Dried calcium sulfate particles were then weighed to calculate the concentration of the suspensions and the concentration was adjusted to account for the concentration of salts contained in the lyophilized DMEM. The described method of calculating concentration of suspension through mass is referred as gravimetric analysis in this paper.

### 2.4 Alginate modification and purification

High molecular weight (180 kDa) alginate (I-1G; Kimica Corporation) was irradiated by a cobalt-60 source to obtain low molecular alginate (25 kDa). For purification, both molecular weights (MW) of alginate were dissolved in DI water, dialyzed (3.5 kDa molecular weight cutoff dialysis membrane; Spectrum Chemicals) for 72 hours, treated with activated charcoal, sterile-filtered, and lyophilized.

For RGD modification of alginate, oligopeptide GGGGRGDSP (Peptide 2.0) was conjugated to alginate using carbodiimide chemistry as described in previously published studies [9]. RGD peptide conjugation chemistry was modified so that the final concentration of RGD peptide in reconstituted 2 wt% hydrogel (for cell culture) was 1.5 mM. RGD modified alginates were then purified using the purification procedure mentioned above.

### 2.5 Hydrogel Fabrication

To fabricate alginate hydrogels for uniaxial compression testing, purified alginates were reconstituted at 2.5 wt% concentration in serum-free DMEM (Life Technologies). Calcium sulfate working solution was prepared at 0.084g/mL concentration in DMEM. Then, to fabricate hydrogel of desired calcium sulfate concentration, two syringes were prepared. One syringe had alginate solution, and the second syringe had DMEM with predetermined amount of calcium sulfate working solution. After removing bubbles inside, the two syringes were connected via Luer lock coupler, rapidly mixed, and the resulting pre-gel solution was quickly cast between two glass plates with 2 mm spacing. The casted hydrogel’s final alginate concentration is 2 wt%. Subsequently, the gels were allowed to form at room temperature for 45 minutes, and then sectioned into disks using a 10 mm biopsy punch. The gel disks were equilibrated for 24 hours at 37 °C in DMEM before compression testing.

### 2.6 Mechanical Test

For compression testing, the gel disks were placed onto the stage of MTS Criterion 43 (MTS). Excess liquid on the gel disks was removed before testing. Then, the compression plate was lowered to just above the gel surface and the test was initiated at a strain rate of 2 mm/min until 15% strain, at which point the strain was held constant and the load was measured until it reached half of its peak value for stress relaxation test. Initial Young’s modulus was calculated by performing linear regression on the initial linear region of the compression test (∼5-10% strain).

### 2.7 3D Encapsulation of NIH-3T3 Cells (3D Culture)

For 3D culture, NIH-3T3 cells were used to obtain a final cell encapsulation density of 5 x 10^6^ cells/mL. RGD-coupled I-1G alginate was first reconstituted at 3.0 wt%. A syringe with cell suspension was used to gently mix cells with alginate in another syringe using Luer lock coupler. Then, the syringe containing cell and alginate mixture was connected via Luer lock coupler to a syringe with calcium sulfate and rapidly mixed, and the resulting pre-gel solution was quickly cast between two glass plates with 1 mm spacing. The casted hydrogel’s final alginate concentration is 2 wt%. Subsequently, the gels were allowed to form at room temperature for 45 minutes, and then sectioned into disks using an 8 mm biopsy punch. The gel disks were cultured in a 24-well plate at 37 °C in 750 μL of DMEM with 10% FBS for a total of 10 days with 50% media exchange on every 3 days.

### 2.8 Viability Assay

On days 0 and 7 of 3D culture, gel disks were stained and imaged for cell viability using calcein-AM and propidium iodide with Hoechst counterstain. First, gel disks were each removed from cell culture media and placed into 400 μL of washing buffer (HEPES supplemented with 2 mM Ca^2+^) in a 24-well plate. A staining solution was prepared with 3 µM Hoechst 33342 (Tocris), 4 µM calcein-AM (eBioscience), and 3 µM propidium iodide (Invitrogen). Then, the washing buffer was aspirated and 400 μL of staining solution was placed into each well. The gel disks were allowed to incubate for 45 min at 37 °C in the staining solution. Next, the staining solution was aspirated and 400 μL washing buffer was placed into each well. The gel disks were allowed to incubate for 15 minutes before another exchange of washing buffer and imaging. For imaging, the gel disks were placed onto a glass slide with 50 μL of washing buffer to keep them hydrated. An EVOS-M5000 microscope was used to take 200-micron Z-stacks of each gel, starting 100 microns away from the bottom surface. ImageJ was used to quantify the max intensity Z-stack projection images for live/dead viability.

### 2.9 F-actin Staining

On day 10 of 3D culture, the gel disks were double washed with HEPES buffer supplemented with 1.5mM Ca^2+^ for 5 minutes each and fixed with 4% paraformaldehyde for 20 minutes at room temperature. The gel disks were then washed twice for 5 minutes and permeabilized for 30 minutes at room temperature using 0.02% Triton X-100 in HEPES buffer supplemented with 1.5 mM Ca^2+^. Permeabilization buffer was then replaced with Phalloidin-iFluor488 (Abcam) solution (1:1000 dilution in HEPES buffer supplemented with 1.5 mM Ca^2+^) and incubated at 4°C overnight. Phalloidin solution was washed and counterstained with Hoechst. For imaging, LSM800 confocal microscope (Zeiss) was used to obtain 50 μm Z-stack images with 3.5 μm sections.

## 3. Results

### 3.1 Calcium Sulfate Microparticle Fabrication and Particle Size Characterization

Different sizes of calcium sulfate microparticles were prepared by mechanical ball-milling of as-received particles and followed by ethanol sterilization (Fig. 1b). After examination of ball-milling parameters (total mass, ball-to-powder mass ratio, milling time, speed), milling time variation was selected as a method of producing microparticles of distinct sizes. Particles sizes measured using optical microscopy showed that as-received calcium sulfate particles had an average size of 27 μm, whereas as-received particles after autoclave sterilization resulted in particles with an average size of 42 μm (Fig. 1c, 1d). Average calcium sulfate particle size after 1 minute, 2 minutes, and 5 minutes of milling was 15 μm, 8.4 μm, and 5.7 μm, respectively (Fig. 1e, 1f). This shows that autoclave treatment increases the size of the as-received particles, while ball milling followed by ethanol treatment decreases the size of particles.

### 3.2 Smaller Calcium Sulfate Microparticles Form More Stable Suspensions

Calcium sulfate suspension stability was measured by the duration of suspension homogeneity and particle sedimentation kinetics. Gravimetric analysis (mass measurement of lyophilized suspension) was used to quantify the suspension concentrations. Calcium sulfate suspension stability was compared for the different size particle samples, and we found that smaller calcium sulfate particles formed more stable suspensions (Fig. 2). 5.7 μm calcium sulfate particles (5-minute milled) were extremely stable in suspension, maintaining a relatively high concentration of suspension even after 2 minutes of sedimentation. 8.4 μm (2-minute milled) particles maintained the next highest concentration of suspension, followed by in the order of 15 μm (1-minute milled) particles, 27 μm (as-received) particles, and 42 μm (autoclaved) particles. Overall, all ball milled calcium sulfate particles showed substantially higher concentration of suspension over time compared to autoclaved calcium sulfate particles, which indicates smaller ball milled particles have higher suspension stability.

**Figure 2.**
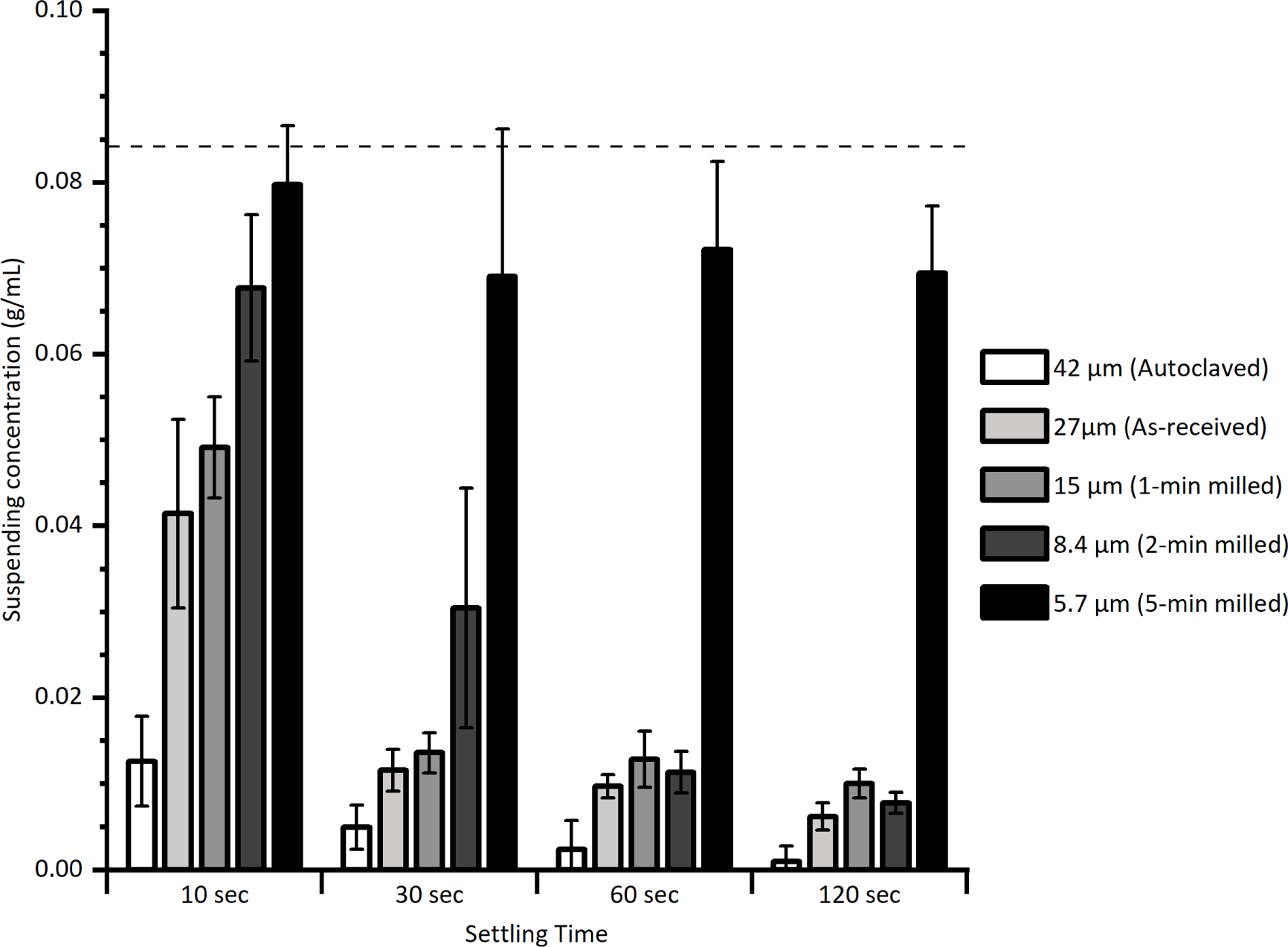
Suspension stability of calcium sulfate particles with different sizes. Suspensions were sampled on different timepoints of 10 sec, 30 sec, 60 sec, 120 sec after stirring, and particle concentration was measured by gravimetric analysis. Particle sedimentation is observed. Dotted line represents the initial concentration of suspension before sedimentation (0.084 g/mL). N = 4-6 for each condition, and error bars represent standard deviation.

### 3.3 Ball-milled Calcium Sulfate Microparticles Form More Optically Homogeneous Gels

Reconstituted low MW alginate (25 kDa) were mixed with calcium sulfate suspension of different particle sizes to be crosslinked into hydrogels (final calcium sulfate crosslinker concentration 43 mM). During gelation, calcium sulfate particles inside the alginate matrix were imaged under brightfield microscope at 20 minutes and 45 minutes after mixing to observe dissolution of particles over time. After hydrogel disks were placed in DMEM for 24 hours of equilibration, additional brightfield microscope images were collected to assess particle dissolution at the end of the equilibration period. Although substantial dissolution for both the 42 μm (autoclaved) and ethanol-sterilized 27 μm (as-received) particles was observed, some large particles or aggregates were visible in the gels at 24 hours at this high calcium sulfate concentration (Fig 3). However, for the smaller milled particle conditions, gels were markedly more homogeneous after only 20 minutes of gelation, and almost no particle aggregates were visible after 24 hours (Fig. 3).

**Figure 3.**
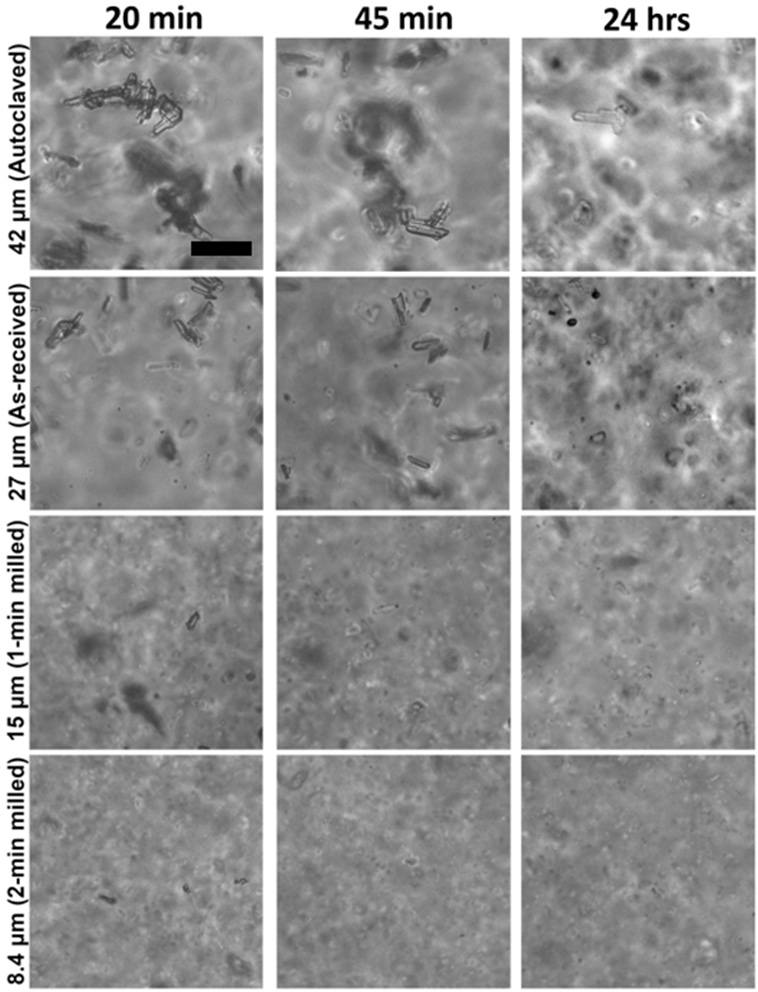
Optical homogeneity of alginate hydrogels (Low MW) crosslinked with different calcium sulfate particles at high concentration (43 mM calcium sulfate). 42 μm (autoclaved), 27 μm (as-received), 15 μm (1-min milled), and 8.4 μm (2-min milled) particles were used to fabricate alginate hydrogels and imaged via optical microscopy at 20 minutes after gel fabrication, 45 minutes after gel fabrication, and after 24 hours of equilibration in DMEM. As particle size decreases, hydrogels became more homogeneous. Scale bar is 50 μm.

### 3.4 Calcium Sulfate Microparticles Can Create Alginate Hydrogels with Tunable Viscoelasticity

Alginate hydrogels of low and high MW and low and high calcium sulfate concentration were fabricated with different particle samples. Each condition was tested using uniaxial compression tests to obtain the initial Young’s modulus and stress-relaxation halftime [9] (the time for the initial stress to be relaxed to half its value during a stress relaxation test).

For high MW alginate, 20 kPa (stiff) and 5 kPa (soft) hydrogels were fabricated by using 22 mM and 9.8 mM of calcium sulfate, respectively. The Young’s modulus of the gels was not affected by varying the type of calcium sulfate particles (Fig. 4a, 4c). The stress-relaxation halftime was also consistent among the 42 μm (autoclaved), 27 μm (as-received), and 15 μm (1-min milled) particle conditions. However, 20 kPa gel crosslinked with 8.4 μm particles (2-min milled) exhibited a lower stress-relaxation halftime. For low MW alginate, 64 mM and 42.7 mM of calcium sulfate were used to form 20 kPa and 5 kPa hydrogels, respectively. Similar to the high MW alginate hydrogels, the Young’s modulus and stress-relaxation halftime of the low MW alginate gels were not affected by varying the type of calcium sulfate particles (Fig. 4b, 4d). However, high concentration (64 mM) of 8.4 μm (2-min milled) calcium sulfate particles caused the alginate to crosslink too quickly, resulting in deformation/fracturing of the hydrogel upon casting. As these samples could not be molded into an intact network using the syringe mixing fabrication method, the 8.4 μm (2-min milled) calcium sulfate condition in the low MW 20 kPa group was not available for Young’s modulus or stress-relaxation test. Overall, the stiffness and stress relaxation of the gels were able to be independently controlled by adjusting the concentration of different calcium sulfate crosslinkers and the MW of alginate, which is consistent with our previous findings [9]. In addition, this was not affected by varying the type of calcium sulfate particles under our testing conditions. Based on the hydrogel formation and mechanical property results, 42 μm (autoclaved) and 15 μm (1 min milled) particles were selected to be investigated further in the following cell culture experiments.

**Figure 4.**
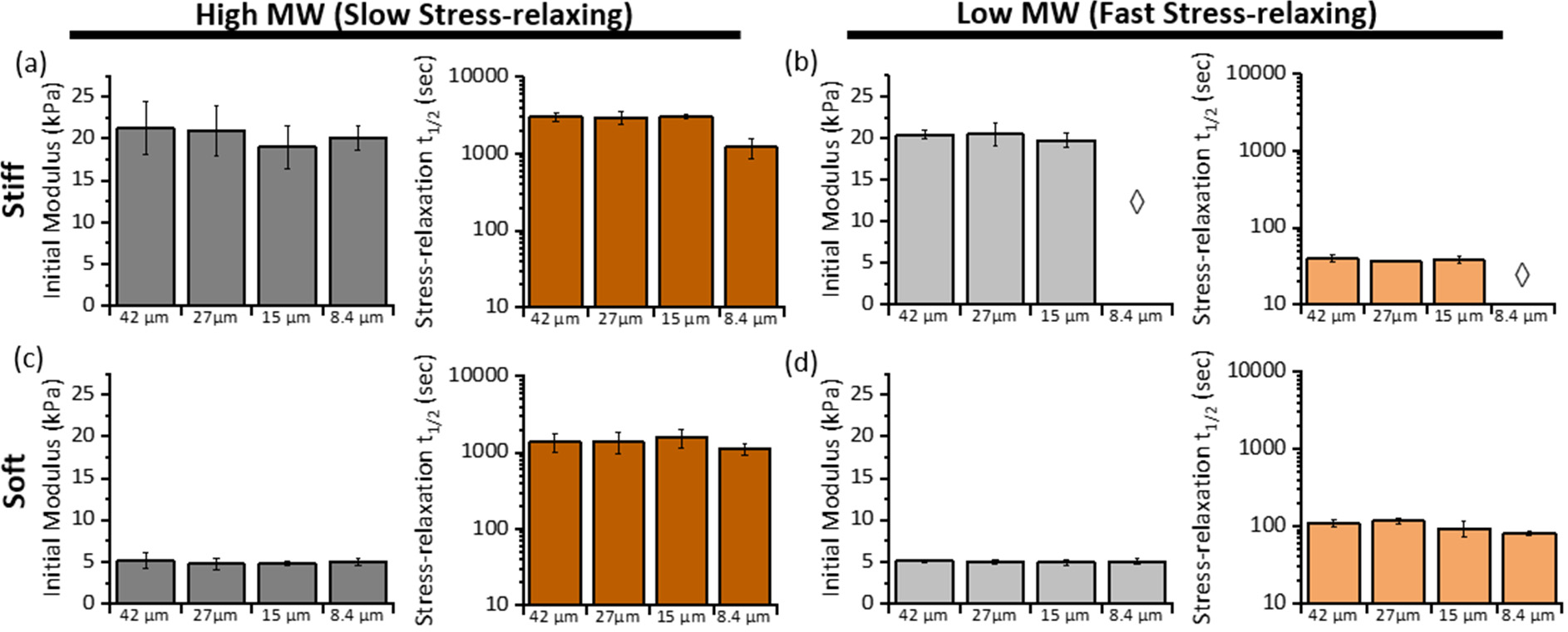
Mechanical properties of alginate hydrogels. Initial Young’s modulus and stress-relaxation halftime (t_1/2_) were measured for (a) stiff, slow-relaxing gels, (b) stiff, fast-relaxing gels, (c) soft, slow-relaxing gels, (d) soft, fast-relaxing hydrogels. Hydrogels were crosslinked using different particles: 42 μm particles (autoclaved), 27 μm particles (as-received), 15 μm particles (1-minute milled), and 8.4 μm particles (2-minute milled). ◊ indicate samples not available for measurement. N = 9 hydrogel disks per group were measured using compression test, and error bars represent standard deviation.

### 3.5 Calcium Sulfate Microparticles Result in Gels with Consistent Quality for 3D Culture

To assess the effect of different calcium sulfate particle size on viscoelastic gels for cell culture, we compared 3D culture of NIH-3T3 fibroblasts results between 42 μm (autoclaved) calcium sulfate particles and 15 μm (1-min milled) calcium sulfate particles as crosslinkers. Using both high and low MW alginate with RGD modification, we encapsulated NIH-3T3 fibroblasts at the density of 5 x 10^6^ cells/mL in 20 kPa and 5 kPa hydrogels with slow or fast stress relaxation properties. Viability of the encapsulated cells was assessed using Calcein-AM and propidium iodide live/dead staining on the day of encapsulation (day 0) and day 7 of 3D culture (Fig. 5). Viability of cells in all conditions was maintained around 80%, indicating that the encapsulation process and the culture environment were suitable for 3D cell culture. Observation of cell morphology in different viscoelastic environments was conducted using F-actin staining of 3D cultures on day 10 (Fig. 6). Slow stress-relaxing alginate hydrogels suppressed cell spreading in both 20 kPa and 5 kPa conditions regardless of the calcium sulfate crosslinker particles. In contrast, the fast stress-relaxing alginate hydrogels resulted in substantial cell spreading in both 20 kPa and 5 kPa conditions for both 42 μm (autoclaved) and 15 μm (1-min milled) calcium sulfate crosslinker particles.

**Figure 5.**
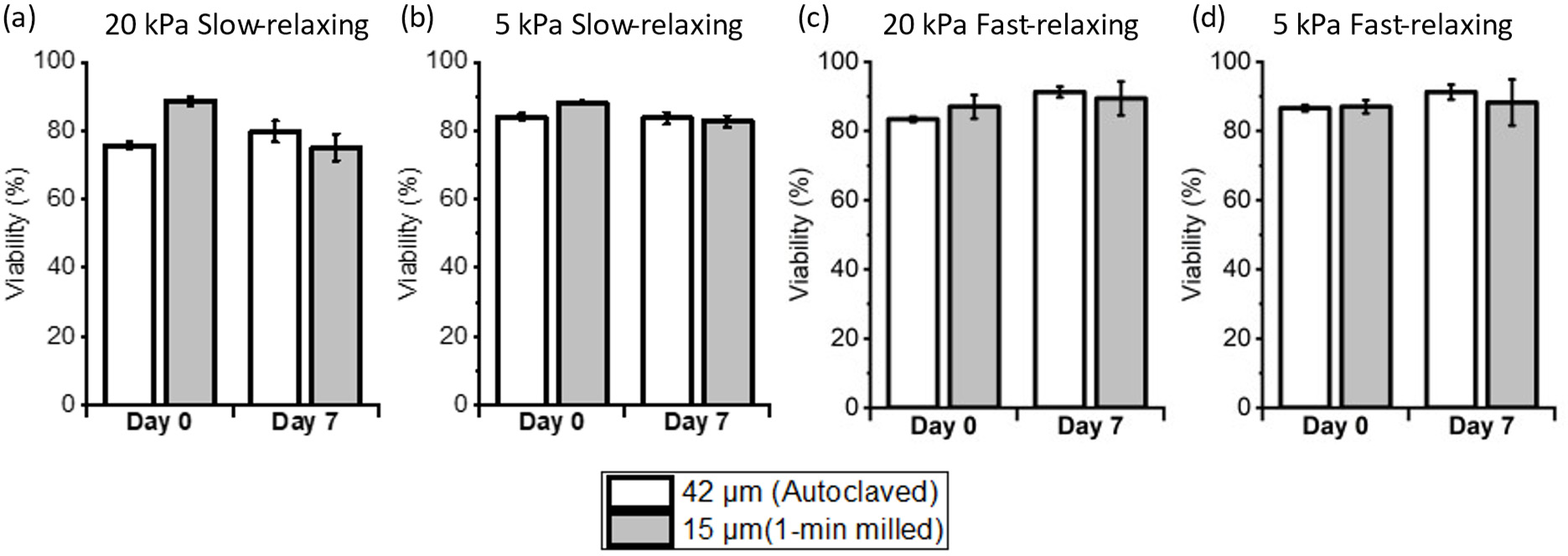
3D culture viability of NIH-3T3 cells in viscoelastic hydrogels crosslinked with 42 μm (autoclaved) or 15 μm (1-min milled) calcium sulfate particles. Viability of cells on day 0 and day 7 of 3D encapsulation in (a) 20 kPa slow-relaxing hydrogel, (b) 5 kPa slow-relaxing hydrogel, (c) 20 kPa fast-relaxing hydrogel, (d) 5 kPa fast-relaxing hydrogel. N=3 per condition; error bars represent standard deviation.

**Figure 6.**
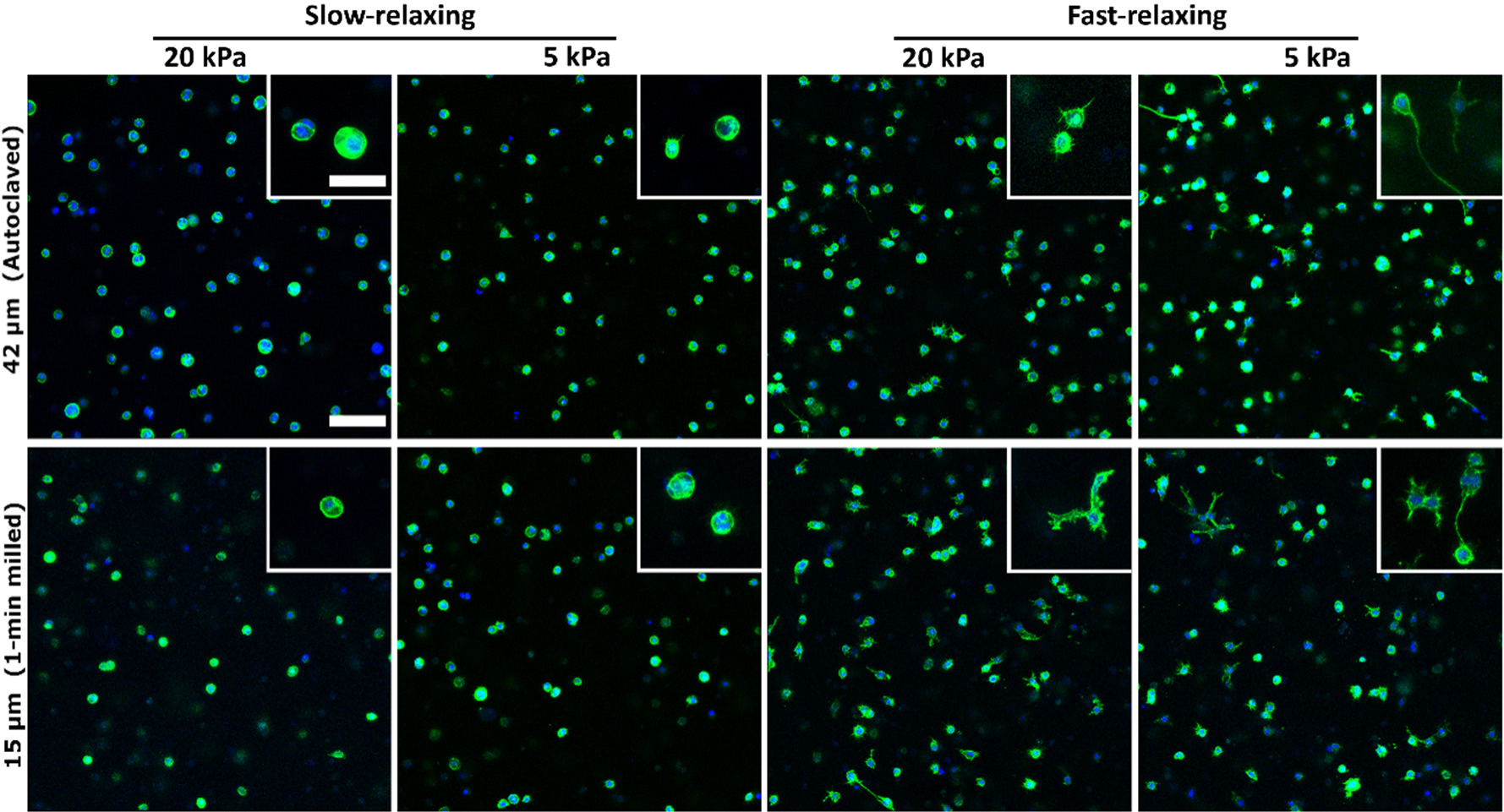
NIH-3T3 cell spreading in 3D culture in alginate gels crosslinked with 42 μm (autoclaved) or 15 μm (1-min milled) calcium sulfate particles. Representative images of 3D cultured NIH-3T3 cells on day 10 of encapsulation. F-actin is green and nucleus is blue. Method of particle preparation did not affect cell spreading result for each stiffness and stress relaxation gel condition. Gels with slow-relaxing property suppress cell spreading regardless of crosslinker particle size, while gels with fast-relaxing property result in substantial cell spreading. Scale bars represents 100 μm for larger images and 50 μm for insets.

## 4. Discussion

We modified the size of calcium sulfate particles to tune the stability of crosslinker suspension for alginate crosslinking. Ball milling of gypsum ore was reported to be a viable method to achieve particle sizes ranging from hundreds of micrometers to a few micrometers [19]. Among a number of ball milling parameters (ball size, ball-to-powder ratio, milling time, etc.), we found that by adjusting the time of the ball milling process, reduced and distinct sizes of calcium sulfate particles for suspension formulations can be easily achieved (Fig. 1f). Particle sterilizations for cell culture applications can be achieved by a number of methods, such as gamma irradiation and autoclaving. However, some of these techniques are not easily accessible and can alter the physical properties of particles (e.g. size, crystallinity, hydration) [20, 21]. We found that autoclaving of calcium sulfate dihydrate particles can increase particle size (Fig. 1f), so 70 % ethanol was used to sterilize the ball milled particles.

As expected, particle size was a critical factor for suspension stability, and calcium sulfate suspension stability was enhanced with reduced particle sizes (Fig. 2). However, there is a fine balance between calcium sulfate particle size and the gelation rate. On the one hand, smaller particles can reduce gel fabrication variability caused by particle sedimentation in the crosslinker suspension, and they also improve gel homogeneity due to their faster dissolution. On the other hand, if the dissolution of small calcium sulfate particles is too fast, it will lead to a high gelation rate, which can prevent adequate mixing of cells and alginate solution with calcium sulfate suspension before the gel fully solidifies. The 5.7 μm (5-minute milled) calcium sulfate particles are such an example. The small particles produced a stable suspension, but they caused instantaneous crosslinking when used to fabricate alginate hydrogels, and subsequent mixing would lead to gel fractures. Therefore, we chose four distinct particle samples to examine hydrogel properties in this study: 42 μm particles (autoclaved), 27 μm particles (as-received), 15 μm particles (1-minute milled), and 8.4 μm particles (2-minute milled).

We have previously found that tuning the molecular weight (MW) of alginate polymer enables to tune the viscoelasticity (e.g. stress-relaxation) of the crosslinked hydrogels [9]. In this work, we utilized 180 kDa and 25 kDa MW alginate polymers to evaluate the effect of calcium sulfate particle size on the mechanical properties of viscoelastic alginate gels. We found that alginate gels crosslinked with 42 μm, 27 μm, or 15 μm calcium sulfate particles had consistent stiffness and stress relaxation properties across these particle sizes, which demonstrates that the gel mechanical properties are not affected by the particles size within this range (Fig. 4). As the 15 μm ball-milled particles exhibit enhanced suspension stability (Fig. 2) and form more homogenous gels at high concentration compared to the larger particles (Fig. 3), they offer significant advantages in fabricating viscoelastic alginate gels. However, when even smaller calcium sulfate particles were used (8.4 μm, 2-minute milled), high concentration of the 8.4 μm particles instantaneously crosslinked low MW alginate due to their fast dissolution (Fig. 4). Overall, the 15 μm size (1-minute milled) particles provide the best balance between suspension stability and gelation rate.

Tunable viscoelastic hydrogels are important tools to study how matrix viscoelasticity influence cell activity, so we also evaluated the effect of calcium sulfate particle size on 3D cell culture. NIH-3T3 fibroblasts were 3D cultured in RGD-coupled alginate hydrogels with different stiffness and stress relaxation properties. Cell viability remained consistent (∼ 80%) in all hydrogels, regardless of whether they were crosslinked using 42 μm (autoclaved) or 15 μm particles (1-minute milled) (Fig. 5). Our previous study found that cell spreading is suppressed in hydrogels with slow stress relaxation properties while cells in fast-relaxing gels exhibit enhanced spreading [9]. Consistent with previous findings, distinct differences in cell spreading were observed between slow-relaxing and fast-relaxing hydrogels regardless of the type of calcium sulfate particles used for crosslinking (Fig. 6). In summary, the viability and cell morphology results show that 15 μm ball milled calcium sulfate particles can serve as highly effective crosslinkers in the fabrication of homogeneous viscoelastic alginate hydrogels for 3D cell encapsulation and culture.

## 5. Conclusion

The 15 μm calcium sulfate particles created using the ball milling method in this study are substantially more stable in suspension than unprocessed or simply autoclaved particles which are commonly used to fabricate viscoelastic alginate hydrogels. The 15 μm particles also result in more homogeneous hydrogels with a balanced gelation rate. They can be used to fabricate alginate hydrogels with tunable viscoelasticity that is similar to previous studies using autoclaved particles. In 3D cell culture, viscoelastic gels fabricated using the 15 μm particles consistently demonstrate the effect of viscoelasticity on cell spreading with high cell viability. Overall, these ball-milled calcium sulfate particles allow for a fabrication method of alginate hydrogels for 3D cell culture that is advantageous over the use of unprocessed or autoclaved particles.

## Acknowledgement

We thank Dr. Tim Weiss for kindly providing access to shaker mill instrument and for helpful discussions. Research reported in this publication was partially supported by the National Institute On Aging of the National Institutes of Health under Award Number R03AG073834. The content is solely the responsibility of the authors and does not necessarily represent the official views of the National Institutes of Health.

## Contribution

JHK and SI formulated and designed the project, performed alginate preparation, calcium sulfate fabrication, hydrogels fabrication, hydrogel testing, imaging, cell culture work, and wrote the manuscript. CT performed suspension stability tests, calcium sulfate fabrication, alginate preparation, hydrogel fabrication, imaging, and cell culture work. SVL designed and performed calcium sulfate fabrication. HK assisted result analysis. LG formulated and designed the project, managed the project, and wrote the manuscript. JHK, SI, CT, SVL, HK and LG read and edited the manuscript.

